# Live Imaging of Intracranial Lymphatics in the Zebrafish

**DOI:** 10.1101/2020.05.13.094581

**Authors:** Daniel Castranova, Bakary Samasa, Marina Venero Galanternik, Hyun Min Jung, Van N. Pham, Brant M. Weinstein

**Author notes:** To whom correspondence should be addressed, Brant M. Weinstein, Section on Vertebrate Organogenesis, Building 6B, Room 4B413, 6 Center Drive, Bethesda, MD 20892.

## Abstract

**Rationale:** The recent discovery of meningeal lymphatics in mammals is reshaping our understanding of fluid homeostasis and cellular waste management in the brain, but visualization and experimental analysis of these vessels is challenging in mammals. Although the optical clarity and experimental advantages of zebrafish have made this an essential model organism for studying lymphatic development, the existence of meningeal lymphatics has not yet been reported in this species.

**Objective:** Examine the intracranial space of larval, juvenile, and adult zebrafish to determine whether and where intracranial lymphatic vessels are present.

**Methods and Results:** Using high-resolution optical imaging of the meninges in living animals, we show that zebrafish possess a meningeal lymphatic network comparable to that found in mammals. We confirm that this network is separate from the blood vascular network and that it drains interstitial fluid from the brain. We document the developmental origins and growth of these vessels into a distinct network separated from the external lymphatics. Finally we show that these vessels contain immune cells and perform live imaging of immune cell trafficking and transmigration in meningeal lymphatics.

**Conclusions:** This discovery establishes the zebrafish as a important new model for experimental analysis of meningeal lymphatic development, and opens up new avenues for probing meningeal lymphatic function in health and disease.

## Introduction

The discovery of meningeal lymphatic vessels in mammals is changing our understanding of how cerebrospinal fluid and interstitial fluid are removed from the brain ^1, 2^. Fluid and macromolecules are collected by lymphatic vessels closely associated with sagittal and transverse sinuses and are drained into the cervical lymph nodes ^2^, and cerebrospinal fluid has been shown to drain through lymphatic vessels lining the base of the skull ^3^. This intracranial lymphatic network is believed to be a route for immune cells to cause neuroinflammation ^4^. Additionally, the role of these lymphatic vessels in draining macromolecules, including amyloid-β, has been shown to be relevant to the development of Alzheimer’s Disease and improving intracranial lymphatic function may be a useful target for disease treatment ^5^. Meningeal lymphatics have also been shown to be a promising new avenue for the treatment of glioblastoma. By increasing VEGF-C (Vascular endothelial growth factor C) levels in the brain, researchers were able to increase the amount of meningeal lymphatic vessels which improved immune surveillance allowing the mouse’s immune system to attack the brain tumor ^6^. The meningeal lymphatic network has great physiological and clinical relevance but has only been described in mammalian research models. Although meningeal lymphatics have been imaged in humans and non-human primates using MRI scans ^7^, the thickness and opacity of the mammalian skull makes high resolution live optical imaging of these vessels extremely challenging, requiring methods such as multi-photon excitation and skull thinning ^4^.

The optical clarity of the zebrafish larva and a number of available transgenic lines for visualizing lymphatic vessels in living animals ^8–11^ have made the zebrafish an excellent model for studying lymphatic development *in vivo*. Use of the zebrafish model has led to important observations regarding the origins and assembly of lymphatic networks, and the genetic pathways that control this ^12–15^, as well as extensive descriptions of larval and juvenile lymphatic networks ^8, 9^. The importance of lymphatic vessels during heart regeneration in zebrafish has also been described ^16, 17^ Close examination of the developing zebrafish brain has led to a thorough description of unusual perivascular lymphatic-related cells called “Fluorescent Granular Perithelial cells” (FGPs) aka “Mato cells” ^18^, “meningeal mural Lymphatic Endothelial Cells” (muLECs) ^19^, or “Brain Lymphatic Endothelial cells” (BLECs) ^20^ with a macrophage-like morphology and a high capacity to take up macromolecules. However, while these cells have a gene expression profile very similar to that of lymphatic endothelial cells they do not form tubes under normal physiological conditions, and *bona fide* intracranial lymphatics have not been described in the zebrafish.

Here, we document a complex intracranial lymphatic vessel network in the juvenile and adult zebrafish comparable to that found in mammals, describe its development from the facial lymphatic vascular plexus, and demonstrate its function in brain fluid clearance and immune cell trafficking. This discovery establishes an important new model for experimental analysis of meningeal lymphatic development and function in health and disease.

## Materials And Methods

### Fish Husbandry and Fish Strains

Fish were housed in a large zebrafish dedicated recirculating aquaculture facility (4 separate 22,000L systems) in 6L and 1.8L tanks. Fry were fed rotifers and adults were fed Gemma Micro 300 (Skretting) once per day. Water quality parameters were routinely measured and appropriate measures were taken to maintain water quality stability (water quality data available upon request). The following transgenic fish lines were used for this study: *Tg(mrc1a:eGFP)^y251^* ^8^, *Tg(−5.2lyve1b:DsRed)^nz101^* ^9^, *Tg(prox1aBAC:KalTA4-4xUAS-E1b:uncTagRFP)^nim5^* ^11^, *Tg(kdrl:mcherry)^y206^* ^21^, *Tg(lyz:DsRed2)^nz50^* ^22^, *Tg(Ola.Sp7:mCherry-Eco.NfsB)^pd46^* previously known as *Tg(osterix:mCherry-NTRo)^pd46^* ^23^. Most of the lines imaged were maintained and imaged in a *casper* (*roy, nacre* double mutant ^24^) genetic background in order to increase clarity for visualization of the meninges by eliminating melanocyte and iridophore cell populations from the top of the head.

### Image Acquisition

Confocal images of intracranial lymphatics were acquired using a Nikon Ti2 inverted microscope with Yokogawa CSU-W1 spinning disk confocal, Hamamatsu Orca Flash 4 v3 camera with the following Nikon objectives: 4X Air 0.2 N.A., 10X Air 0.45 N.A., 20X water immersion 0.95 N.A., and 25X silicone immersion 1.05 NA, 40X water immersion 1.15 NA, 60X water immersion 1.20 N.A. Stereo microscope pictures were taken using a Leica M205 microscope using MultiFocus focus stacking. The large size of the juvenile and adult zebrafish head often required tile acquisitions that were later stitched using Nikon Elements software.

### Image Processing

Images were processed using Nikon Elements and Photoshop. *Tg(mrc1a:egfp)^y251^* expression is significantly stronger in FGPs than in intracranial lymphatics. To account for this brightness discrepancy, in some images where intracranial lymphatics are displayed as a different color than FGP’s, the brightness of FGP’s and intracranial lymphatics was adjusted separately to make the intracranial lymphatics easier to see. We determined which vessels were inside vs. outside of the skull by scrolling through z stack planes of images with either *Tg(Ola.Sp7:mCherry-Eco.NfsB)^pd46^* expression or 405 nm autofluorescence of calcified skull bone. Unless otherwise specified, maximum intensity projections of confocal stacks are shown. Focus stacking of confocal images with DIC was done using Nikon Elements EDF (Extended Depth of Focus). 3D rotation movies and time-lapse movies were made using Nikon Elements and exported to Adobe Premiere Pro CC 2019. Adobe Premiere Pro CC 2019 and Adobe Photoshop CC 2019 were used to add labels and arrows to movies and to add coloring or pseudo-coloring. Schematics where made using Adobe Photoshop CC 2019 and Bio Render software.

### Developmental Series

Fish from 14 dpf to 37 dpf were imaged repeatedly by anesthetizing them in 168 mg/L tricaine in system water and mounting them in 0.8% low melting point agarose in a 35mm glass bottomed petri dish (MatTek # P35-1.5-14-C) with the top of their head touching the glass cover slip. After imaging (10-15 minutes) fish were freed from the agarose and revived by placing them back into water from the main aquarium system and applying gentle water flow over the fish using a transfer pipette. Revival sometimes took as long as 30 minutes.

### Juvenile and Adult Intracranial Imaging

Mounting fish older than 30 dpf in low melting point agarose can cause morbidity and death, possibly because of the agarose covering the adult gills. Instead, older fish were mounted by anesthetizing them in 126-168 mg/L tricaine in system water and then placed into a slit in a sponge (Jaece Identi-Plugs L800-D) moistened with tricaine water, cut into a rectangle to fit inside a single chamber imaging dish (Lab-TekII #155360). The sponge containing the fish is placed into the imaging dish with the head against the glass (fish upside down) and the chamber is filled with tricaine water. For longer term imaging (greater than one hour), fish were intubated using a modification of the methods described by C. Xu et al. ^25^ adapted for inverted confocal microscopy (**Supp. Fig. 1 E**). Fish lengths are reported as standard length from the tip of the snout to the base of the tail.

### Angiography and Lymphatic Drainage Injections

Injections were performed as described by Yaniv et al. ^26^. All injections were performed using a Drummond Nanoject II microinjector (Item# 3-000-204) with pulled glass capillary needles (Drummond item # 3-00-203-G/X). One or two injection boluses were given at each injection site with a volume setting of 36.8 nL. Angiography and lymphatic drainage assays were done using undiluted (2 uM) Qtracker™ 705 Vascular labels (Invitrogen cat# Q21061MP). Lymphatic drainage assays were also done using 10,000 MW Cascade Blue™ Dextran (Invitrogen Cat# D1976).

### Alizarin Red Staining

Alizarin red has been shown to be an effective bone stain in living zebrafish ^27^. Adult zebrafish were bathed in 1.9 mM alizarin red bath (400 ul of Alizarin-Red Staining Solution (Sigma Aldrich TMS-008-C 40 mM) in 80 mL of system water) for 40 minutes. After the bath fish were given a one hour rinse in clean system water before imaging.

### Imaging Neutrophils and Intracranial Lymphatics

Living *casper* mutant *Tg(mrc1a:egfp)^y251^*, *Tg(lyz:DsRed2)^nz50^* double-transgenic zebrafish were mounted and imaged as described in the Juvenile and Adult Intracranial Imaging section. The number of neutrophils seen in the cranial region was highly variable from fish to fish. Oxazalone has been used to induce an immune response in zebrafish ^28^. By soaking adult zebrafish in a 0.0005% Oxazolone (4-Ethoxymethylene-2-phenyl-2-oxazolin-5-one, Sigma E0753-5G) solution in system water for 3 hours the number of neutrophils in the cranial region could be greatly increased. This technique was used on the fish shown in **Fig. 7 B-D**.

## Results

### Meningeal lymphatics in the adult zebrafish

Historically, lymphatic vessels have been thought to be excluded from the central nervous system, but several recent reports have documented the presence of lymphatic vessels in the meninges surrounding the mouse brain, particularly adjacent to dural sinuses, and have demonstrated the important function of these vessels for draining cerebrospinal fluid (CSF), their strong association with immune cells, and their importance for immune surveillance of the brain ^1, 2^. Since the zebrafish provides a superb model for *in vivo* analysis of vascular development and function ^29^ and since it has a lymphatic vascular system comparable in most respects to that of mammals ^30, 31^, we sought to determine whether zebrafish also possess intracranial lymphatics, and whether fish could provide a valuable model for live imaging of intracranial lymphatic development and function.

The brain can be visualized through the top of the skull in dorsal views of the heads of adult *casper* (*roy, nacre* double mutant) animals that lack melanin pigment except in their eyes (**Fig. 1A**). The large prominent optic tecta are readily visible in dorsal views of *casper* heads, as are the forebrain and cerebellum (**Fig. 1A,B**). Using a *Tg(mrc1a:egfp)^y251^* zebrafish transgenic reporter line we recently developed for visualizing lymphatic vessels *in vivo* ^8^, confocal imaging of the dorsal heads of living (**Supp Fig. 1A-C,E**) *casper*, *Tg(mrc1a:egfp)^y251^* transgenic adult animals reveals complex networks of superficial lymphatic vessels covering much of the surface of the head, especially over the forebrain area (**Supp Fig. 1A-C**). However, in addition to the superficial lymphatics outside the skull, we also observe an elaborate network of intracranial meningeal lymphatics immediately beneath the skull, particularly over the optic tecta and cerebellum (**Fig. 1B,C, Supp Fig. 1A,B,D**).

**Fig. 1.**
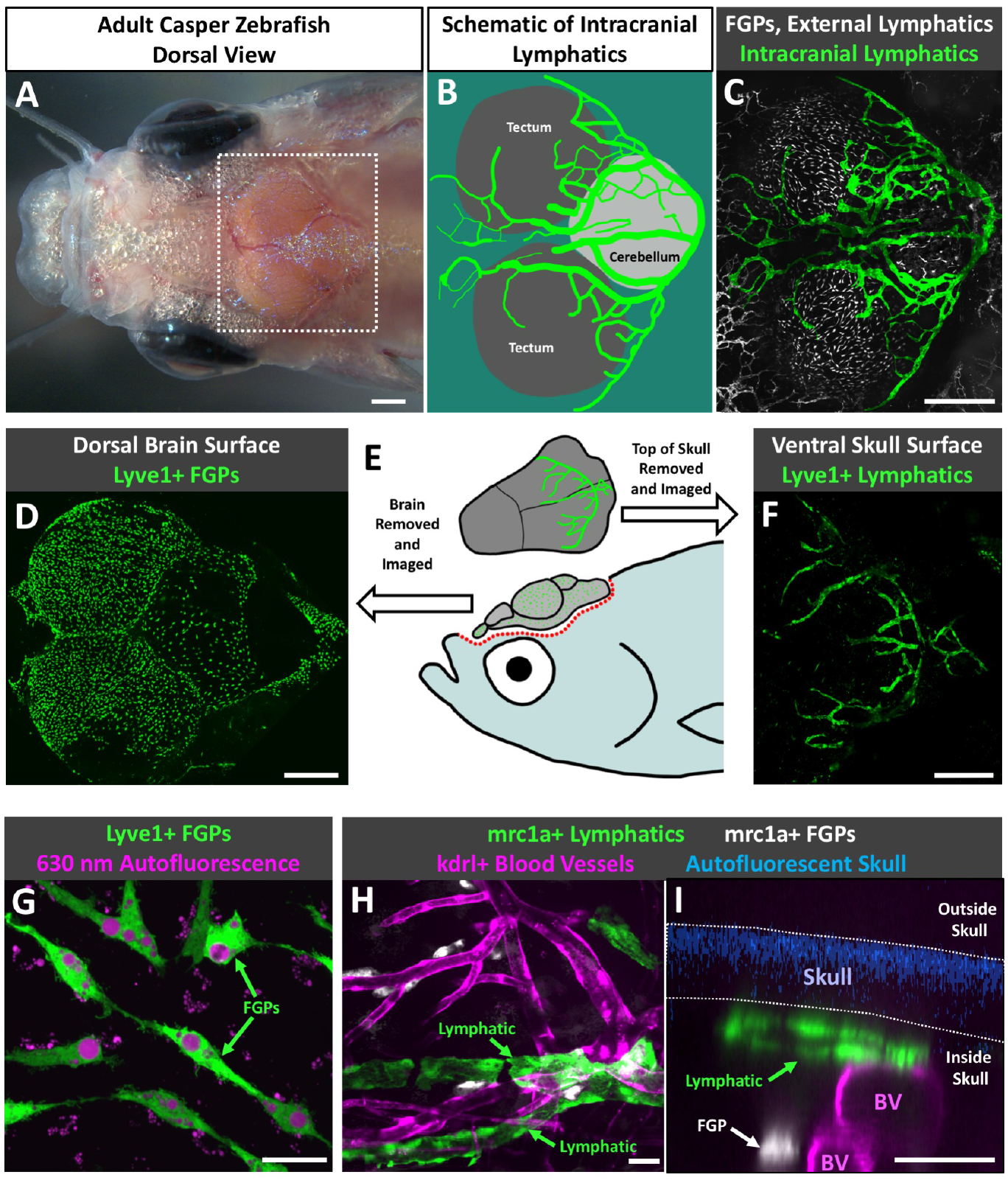
Intracranial lymphatic vessels in the adult zebrafish. **A.** Dorsal view image of the head of an adult *casper (roy, nacre* double mutant) zebrafish. **B.** Schematic diagram of the boxed region in panel A, showing the optic tecta and cerebellum with a typical network of intracranial lymphatic vessels in a young adult zebrafish. **C.** Confocal image of the dorsal head of a *Tg(mrc1a:egfp)^y251^*, *casper* adult zebrafish, with mrc1a+FGPs and superficial lymphatics in grey and intracranial meningeal lymphatics pseudocolored green. **D.** Schematic diagram of dissection of an adult zebrafish head for imaging the dorsal surface of the brain and the ventral surface of the skull. **E.** Confocal image of the dorsal surface of a dissected brain from a *Tg(−5.2lyve1b:DsRed)^nz101^*, *casper* adult zebrafish, with lyve1+ FGPs but no lymphatic vessels. **F.** Confocal image of the ventral (inner) surface of a dissected brain from a *Tg(−5.2lyve1b:DsRed)^nz101^*, *casper* adult zebrafish, with lyve1+ lymphatic vessels but no FGPs. **G.** Higher magnification confocal image of the dorsal surface of a dissected brain removed from a *Tg(−5.2lyve1b:DsRed)^nz101^*, *casper* adult zebrafish, showing individual separated lyve1+ FGPs with characteristic large autofluorescent internal vacuoles. **H.** Higher magnification confocal image of the ventral (inner) surface of a dissected skull removed from a *Tg(−5.2lyve1b:DsRed)^nz101^*, *casper* adult zebrafish, showing lyve1+ lymphatic vessels. **I.** Higher magnification confocal image of the dorsal head of a *Tg(mrc1a:egfp)^y251^, Tg(kdrl:mcherry) ^y206^*, *casper* adult zebrafish. This orthogonal view shows a cross section of an mrc1a+ lymphatic vessel immediately below the blue autofluorescent skull and an mrc1a+ FGP in a deeper layer immediately adjacent to a kdrl+ blood vessel. Unless otherwise noted all images are dorsal views, rostral to the left. Scale bars: 500 um (A,C,E,F) 25 um (G,H,I).

These intracranial lymphatics are distinct and separate from mrc1a-positive “Fluorescent Granular Perithelial” cells (FGPs, aka “Mato Cells,” “MuLECs,” or “BLECs,”), non-tube forming meningeal perivascular cells we and others recently described that are molecularly very similar to lymphatic endothelial cells ^18–20^. FGPs are found in the leptomeninges closely associated with the outer surface of the brain ^18^, and they are readily observed on dissected brains removed from *Tg(−5.2lyve1b:DsRed)^nz101^* transgenic adult animals (**Fig. 1D,E**). However, no lyve1+ lymphatic vessels are observed on the surface of dissected brains. In contrast, the inner surface of dissected skull caps removed from *Tg(−5.2lyve1b:DsRed)^nz101^* transgenic adult animals completely lacks lyve1+ FGPs, and instead shows robust networks of lyve1+ lymphatic vessels (**Fig. 1E,F**). This is analogous to observations from mice, where intracranial lymphatics are found exclusively in the skull-associated meninges and not in the brain-associated leptomeninges ^1, 2^. Closer examination of lyve1+ cells associated with the outer surface of the brain (**Fig. 1D**) and inner surface of the skull (**Fig. 1F**) confirms the distinct identities of the cells in these two locations. The outer surface of the dissected brain contains exclusively FGPs; individual, separated cells immediately adjacent to blood vessels with a macrophage-like morphology and abundant large autofluorescent intracellular vacuoles (**Fig. 1G**). The ventral surface of the skull cap has lyve1+ tubular vessels (**Fig. 1F**) but no FGPs. Orthogonal views of confocal images taken through the intact head of a *casper*, *Tg(mrc1a:egfp)^y251^, Tg(kdrl:mcherry^y206^* double transgenic adult animal confirm that mrc1a+ lymphatic tubes are closely apposed to the skull while mrc1a+ FGPs lie immediately adjacent to kdrl+ blood vessels in a deeper layer (**Fig. 1I, Supp. Movie 1**). Endogenous autofluorescence of calcified bone under ultraviolet light ^32^ allows us to determine which lymphatic vessels are located inside the skull and which are located outside the skull in adult zebrafish without the use of additional transgenics labeling bone, or bone stains (**Supp Fig. 2, Supp Movie 2, Fig 1I**).

To confirm their lymphatic molecular identity, we carried out additional confocal imaging of intracranial lymphatic vessels in *casper*, *Tg(mrc1a:egfp)^y251^*, *Tg(−5.2lyve1b:DsRed)^nz101^* (**Fig. 2A-C**) or *casper*, *Tg(mrc1a:egfp)^y251^, Tg(prox1aBAC:KalTA4-4xUAS-E1b:uncTagRFP)^nim5^* (**Fig. 2D-F**) adult zebrafish to determine whether these different characteristic lymphatic transgenes colocalize in presumptive zebrafish intracranial lymphatics. *mrc1a, lyve1, and prox1a* are widely used markers for lymphatic vessels, and *Tg(mrc1a:egfp)^y251^, Tg(−5.2lyve1b:DsRed)^nz101^*, and *Tg(prox1aBAC:KalTA4-4xUAS-E1b:uncTagRFP)^nim5^* transgenic lines have been developed using the promoters from each of these genes to drive fluorescent reporter expression for live imaging of lymphatic vessels in the zebrafish ^8, 9, 11^. As expected, zebrafish intracranial lymphatic vessels co-express *mrc1a* and *lyve1* (**Fig. 2A-C**) and co-express *mrc1a* and *prox1a* (**Fig. 2D-F**), showing they possess the molecular signature of lymphatic vessels. To further confirm that these vessels are *bona fide* lymphatic vessels and validate both their separation from the blood vessels of the cardiovascular system and their function in intracranial drainage, we carried out blood vessel labeling (via angiography, injection into the dorsal aorta; **Fig. 3A**) and lymphatic vessel labeling (via cranial intracisternal injection; **Fig. 3B** and spinal intracisternal injection; **Supp Fig. 3A**) using Qdot705, blue dextran, or both together to label either blood vessels (**Fig. 3C-E**) or draining intracranial lymphatics (**Fig. 3F-K, Supp Fig. 3C,D**), respectively. Angiographic injection of Qdot705 into the trunk dorsal aorta leads to rapid labeling of the entire circulatory system including intracranial blood vessels without any labeling of adjacent mrc1a:egfp+ intracranial lymphatics (**Fig. 3C-E**). In contrast, intracisternal injection of Qdot705 and blue dextran leads to filling of the brain ventricles, subarachnoid, and subdural spaces (**Supp Fig. 3B**) quickly followed by drainage through and labelling of nearby mrc1a:egfp+ intracranial lymphatics without labelling of adjacent blood vessels (**Fig. 3F-K, Supp Fig. 3C,D, Supp Movie 3**).

**Fig. 2.**
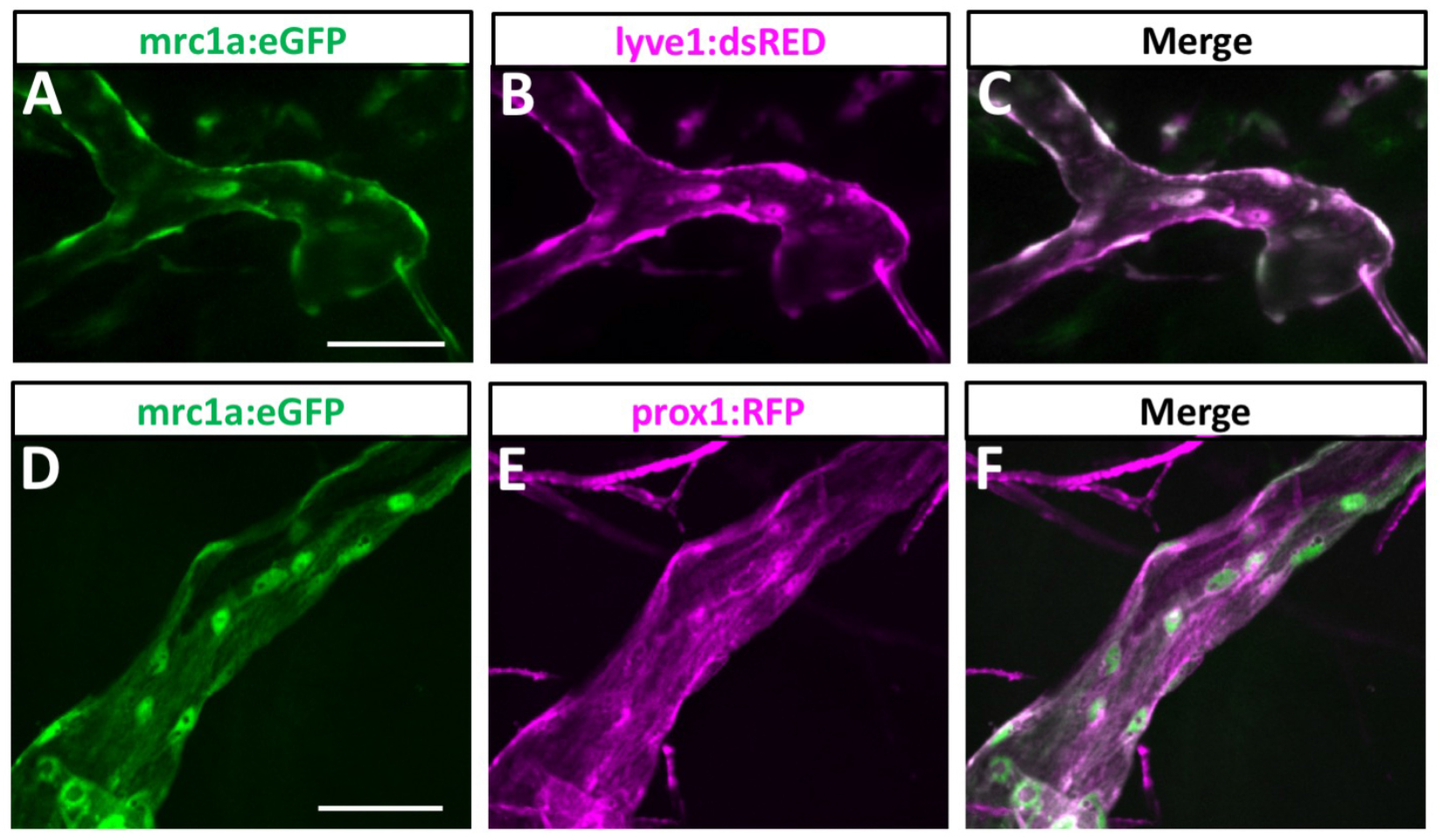
Molecular validation of zebrafish intracranial lymphatics. **A-C.** Confocal imaging of an intracranial lymphatic vessel in the dorsal head of an adult *casper, Tg(mrc1a:egfp)^y251^, Tg(−5.2lyve1b:DsRed)^nz101^* double transgenic zebrafish, showing mrc1a:egfp (A), lyve1b:dsred (B), and combined mrc1a:egfp and lyve1b:dsred (C) images. **D-F.** Confocal imaging of an intracranial lymphatic vessel in the dorsal head of an adult *casper, Tg(mrc1a:egfp)^y251^, Tg(prox1aBAC:KalTA4-4xUAS-E1b:uncTagRFP)^nim5^* double transgenic zebrafish, showing mrc1a:egfp (D), prox1:rfp (E), and combined mrc1a:egfp and prox1:rfp (F) images. Scale bars: 50 um.

**Fig. 3.**
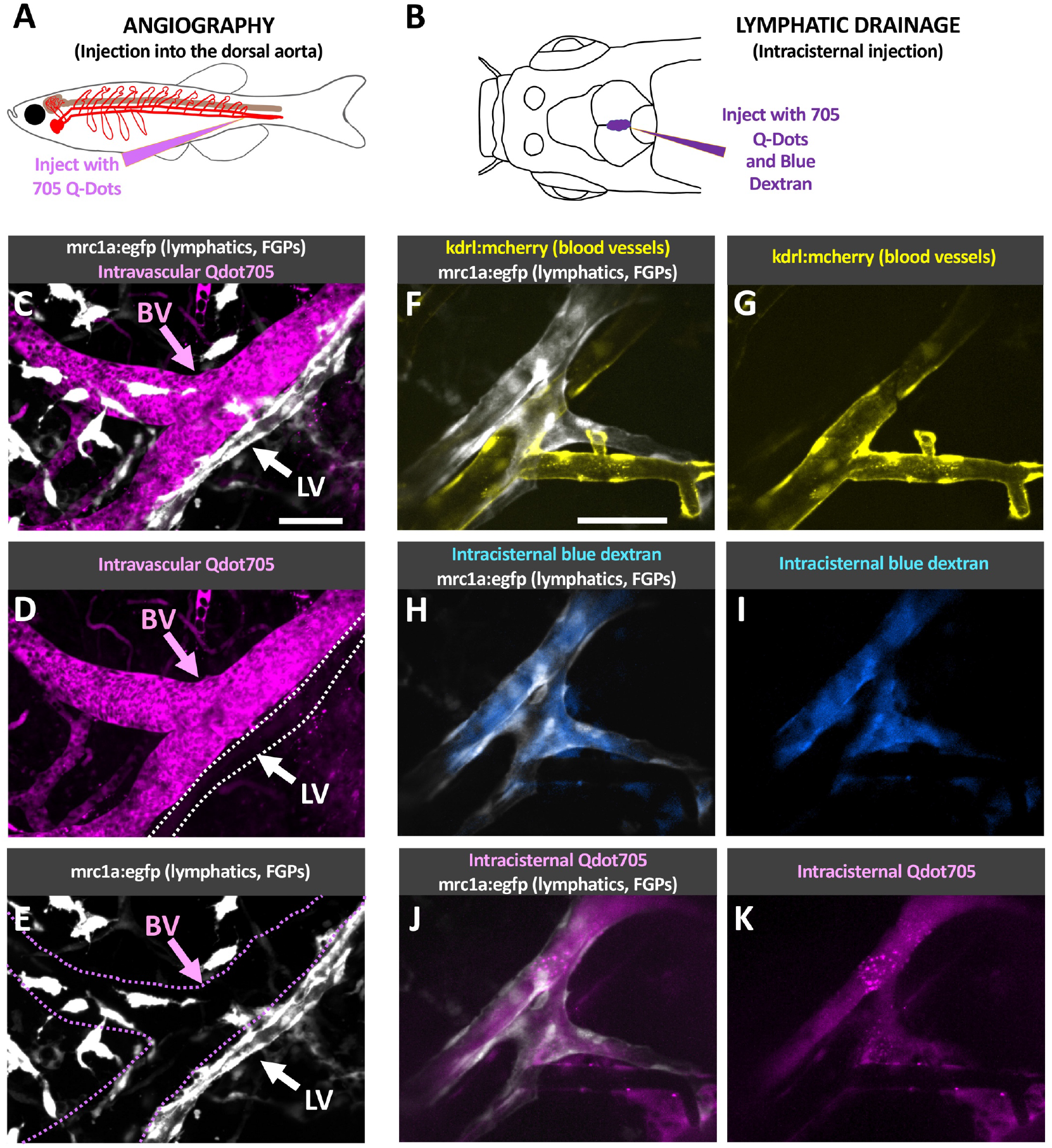
Functional validation of zebrafish intracranial lymphatics. **A.** Schematic diagram of the intravascular injection procedure for filling blood vessels. **B.** Schematic diagram of the intracisternal injection procedure for filling intracranial lymphatics. **C-E.** Confocal images of intracranial blood vessels (BV) and lymphatic vessels (LV) in the dorsal head of a *Tg(mrc1a:egfp)^y251^* adult fish injected intravascularly with Qdot705, showing mrc1a:egfp+ lymphatics and FGPs + Qdot705 (C), Qdot705 alone (D), or mrc1a:egfp+ alone (E). **F-K**. Confocal images of intracranial blood vessels (BV) and lymphatic vessels (LV) in the dorsal head of a *Tg(mrc1a:egfp)^y251^, Tg(kdrl:mcherry) ^y206^* adult fish injected intracranially with Qdot705 and blue dextran, showing mrc1a:egfp+ lymphatics and kdrl:mcherry+ blood vessels (F), kdrl:mcherry+ blood vessels (G), mrc1a:egfp+ lymphatics and blue dextran (H), blue dextran (I), mrc1a:egfp+ lymphatics and Qdot705 (J), Qdot705 (K). Scale bars: 50 um. See **Supp. Movie 3** for 3D renderings and real-time imaging of the vessels shown in panels F-K.

Together these results demonstrate that zebrafish possess a network of intracranial meningeal lymphatic vessels comparable to those described in mammals ^1, 2^.

### Development of intracranial lymphatics in the zebrafish

Having established the existence of intracranial meningeal lymphatics in the adult zebrafish, we sought to understand when and how this network of vessels develops by confocal imaging of the intact dorsal heads of *casper Tg(mrc1a:egfp)^y251^* transgenic zebrafish. Meningeal lymphatics initially sprout from dorsal-medial branches of the bilateral facial lymphatic networks beginning at approximately 9-10 days post fertilization (dpf) or when they are approximately 4 mm in length (**Fig. 4A,E,I, Supp. Movie 4**), initially growing rostrally and laterally along the caudal margin of the cerebellum and optic tecta (**Fig. 4B,F,J, Supp. Movie 4**). These initial branches continue to develop into enlarged sacs and also grow medially eventually connecting across the midline at the caudal edge of the cerebellum (**Fig. 4C,D,G,H,K,L, Supp. Movie 4**). The developing meningeal lymphatics are initially connected to the facial lymphatics from which they emerge on either side (**Fig. 4A,E,I, Supp. Movie 4**), but at later stages of development they become disconnected from the facial lymphatics (**Fig. 4B,C,F,G,J,K, Supp. Movie 4**) connecting instead to more dorsal superficial networks of lymphatic vessels over the cerebellum and hindbrain (**Fig. 4D,H,L, Supp. Movie 4**). The facial lymphatics also connect to this dorsal superficial network at later stages so the developing intracranial lymphatics thus appear to be effectively (albeit indirectly) reconnected to the facial lymphatics in many cases, at least temporarily, although these connections are also lost over time. Although the timing is somewhat variable, the sequence of events in development of the primary intracranial network is quite stereotypical, as shown by imaging of numerous different animals from 14 dpf to 36 dpf (**Supp. Fig. 4**).

**Fig. 4.**
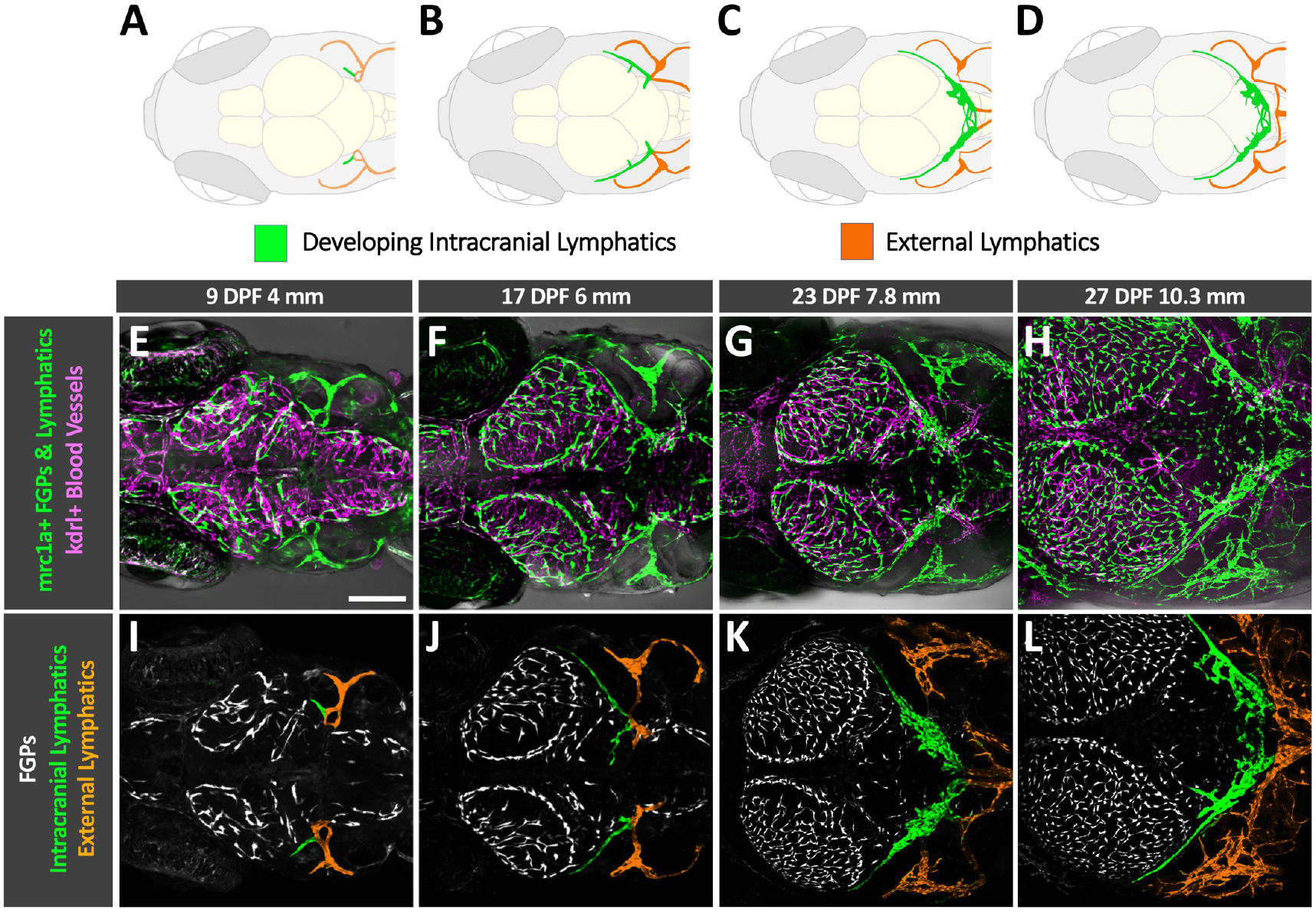
Initial development of intracranial lymphatics in larval zebrafish. **A-D.** Schematic diagrams showing brain structures, developing intracranial lymphatics (green), and external lymphatics (orange) in the dorsal heads of 9 dpf/4 mm (A), 17 dpf/6 mm (B), 23 dpf/7.8 mm (C), and 27 dpf/10.3 mm (D) zebrafish. Each diagram illustrates features in the corresponding confocal micrograph below (panels E-H). OT, optic tecta; C, cerebellum. **E-H.** Confocal images of kdrl:mcherry positive blood vessels (magenta) and mrc1a:egfp positive lymphatic vessels and FGPs (green) in the dorsal heads of 9 dpf/4 mm (E), 17 dpf/6 mm (F), 23 dpf/7.8 mm (G), and 27 dpf/10.3 mm (H) *Tg(mrc1a:egfp)^y251^, kdrl:mcherry)* double-transgenic zebrafish. **I-L.** Colored versions of the same confocal image fields shown in panels E-H with only the mrc1a:egfp fluorescence channel visible, revealing FGPs (white, uncolored), external lymphatics (colored orange), and developing intracranial lymphatics (colored green). Scale bar: 250 um. See Supp. Movie 4 for 3D renderings of the same image stacks shown in this figure.

Although the initial development of the future intracranial lymphatics precedes skull formation, continued later development of intracranial lymphatics occurs concomitantly with growth and closure of the bony skull plates, with growth of the plates across the top of the head successively covering consecutive portions of the meningeal lymphatics. We used confocal imaging of the intact dorsal heads of *casper*, *Tg(mrc1a:egfp)^y251^, Tg(Ola.Sp7:mCherry-Eco.NfsB)^pd46^* (previously known as *Tg(osterix:mCherry-NTRo)^pd46^)* double transgenic zebrafish to simultaneously image Mrc1a:egfp positive developing lymphatics and sp7:mCherry+ developing skull plate bone. Sp7+ skull plates begin to appear at approximately 13 dpf/6.2 mm at the lateral corners of the dorsal head (**Fig. 5A**, arrows). The plates grow continuously over approximately the first month of development, gradually closing the bone-free space at the top of the head and successively covering developing intracranial lymphatics (diagrammed in **Fig. 5B**). By approximately 17 dpf/8.0 mm the growing plates abut portions of the developing future intracranial lymphatics along the caudal margin of the cerebellum (**Fig. 5C-E**, arrows in D,E, **Supp Movie 5**). As seen in a 19 dpf/8.9 mm animal, the growing plates of the optic tecta begin to cover portions of the developing intracranial lymphatics (**Fig. 5B,F-H**, arrows in G,H, **Supp Movie 6**). The mrc1a+ intracranial lymphatics are further covered by the growing sp7+ plates in a 26 dpf/11.7 mm animal (**Fig. 5I-K**, arrows in J,K), and are nearly completely covered by the fused skull plates in a 35 dpf/12.0 mm animal (**Fig. 5L-N**). Higher magnification images of intracranial lymphatics in the 35 dpf/12.0 mm animal confirm that mrc1a+ vessels are immediately below the sp7+ skull (**Fig. 5M,N**).

**Fig. 5.**
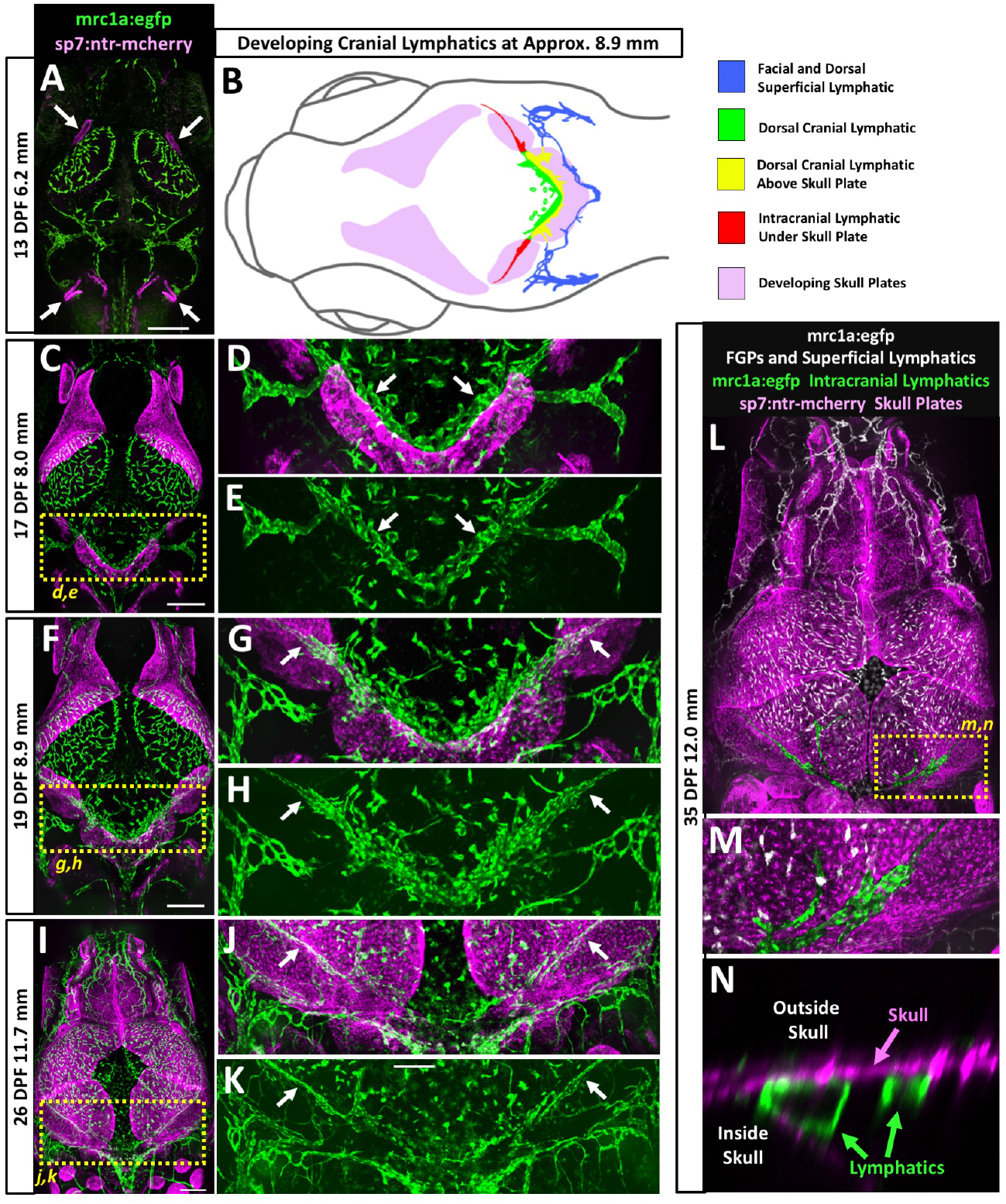
Intracranial lymphatic development and skull formation. **A,C,F,I,L.** Overview confocal images of mrc1a:egfp+ lymphatic vessels and FGPs (green) and sp7:mcherry+ developing skull plates (magenta) in the dorsal heads of 13 dpf/6.2 mm (A), 17 dpf/8.0 mm (C), 19 dpf/8.9 mm (F), 26 dpf/11.7 mm (I), and 35 dpf/12.0 mm (L) *Tg(mrc1a:egfp)^y251^, Tg(Ola.Sp7:mCherry-Eco.NfsB)^pd46^* double-transgenic zebrafish. Arrows in panel A note beginning of skull plate formation. Yellow dashed boxes in panels C, F, I, and L note the areas shown in the corresponding higher magnification images in the panels to the right (D,E,G,H,J,K) or below (M,N). **B.** Schematic diagram corresponding to the confocal image of a 19 dpf/8.9 mm *Tg(mrc1a:egfp)^y251^, Tg(Ola.Sp7:mCherry-Eco.NfsB)^pd46^* double-transgenic zebrafish shown in panel F. Diagram shows developing skull plates (pink), superficial lymphatics (blue), still-uncovered intracranial lymphatic (green), portions of intracranial lymphatic network present above edge of growing skull plates (yellow), and portions of intracranial lymphatics already covered by the growing skull plates (red). **D,G,J.** Higher magnification confocal images of the yellow boxed regions in the panels to the left, showing mrc1a:egfp+ lymphatics (green) and sp7:mcherry+ developing skull plates (magenta). White arrows note lymphatics adjacent to (D) or under (G,J) skull plates. **E,H,K.** Higher magnification confocal images of the yellow boxed regions in the panels to the left, showing only mrc1a:egfp+ lymphatics (green). White arrows note lymphatics adjacent to (E) or under (H,K) skull plates. **M.** Higher magnification confocal image of the yellow boxed region in panel L, showing mrc1a:egfp+ meningeal lymphatics (green) below the nearly completed sp7:mcherry+ skull (magenta). **N.** Higher magnification orthogonal (side) view confocal image of the yellow boxed region in panel L, showing mrc1a:egfp+ meningeal lymphatic tubes (green) inside the skull below the sp7:mcherry+ skull (magenta). Scale bars: 250 um.

From approximately 3 weeks of development the initial relatively simple larval network of intracranial lymphatic vessels found along the caudal margin of the optic tecta and cerebellum (**Fig. 4C,G,K**) grows rostrally and elaborates into a much more complex juvenile and adult intracranial lymphatic vascular system, as visualized by confocal imaging of the intact dorsal heads of *casper Tg(mrc1a:egfp)^y251^* transgenic zebrafish (**Fig. 6A-D**). In a 30 dpf/16 mm animal rostral projections are observed just beginning to emerge from the larval intracranial lymphatics (**Fig. 6A**), while in a slightly more developed 40 dpf/17 mm animal the rostral projections are extended branches curving around the cerebellum toward the dorsal midline (**Fig. 6B**). In a 41 dpf/19 mm animal the lymphatics form a network circumscribing and also beginning to cross the dorsal surface of the cerebellum, and also extending rostrally between the optic tecta on either side of the midline dural sinus (**Fig. 6C**). An even more elaborate network can be seen in a 45 dpf/24 mm animal, with intracranial vessels further extending across the optic tecta and elsewhere (**Fig. 6D**). Although the complete network of intracranial lymphatics become difficult to image through the skull after the first 1-1/2 months of development, the intracranial lymphatics in older animals appear very similar to the network observed in 1-1/2 month old animals, albeit with somewhat increasing complexity (data not shown). Higher magnification imaging of developing juvenile lymphatics suggests that the elaborate rostral lymphatics grow via both sprouting lymphangiogenesis and formation and fusion of locally developing small lymphatic cell clusters/sacs(**fig. 6E-H**).

**Fig. 6.**
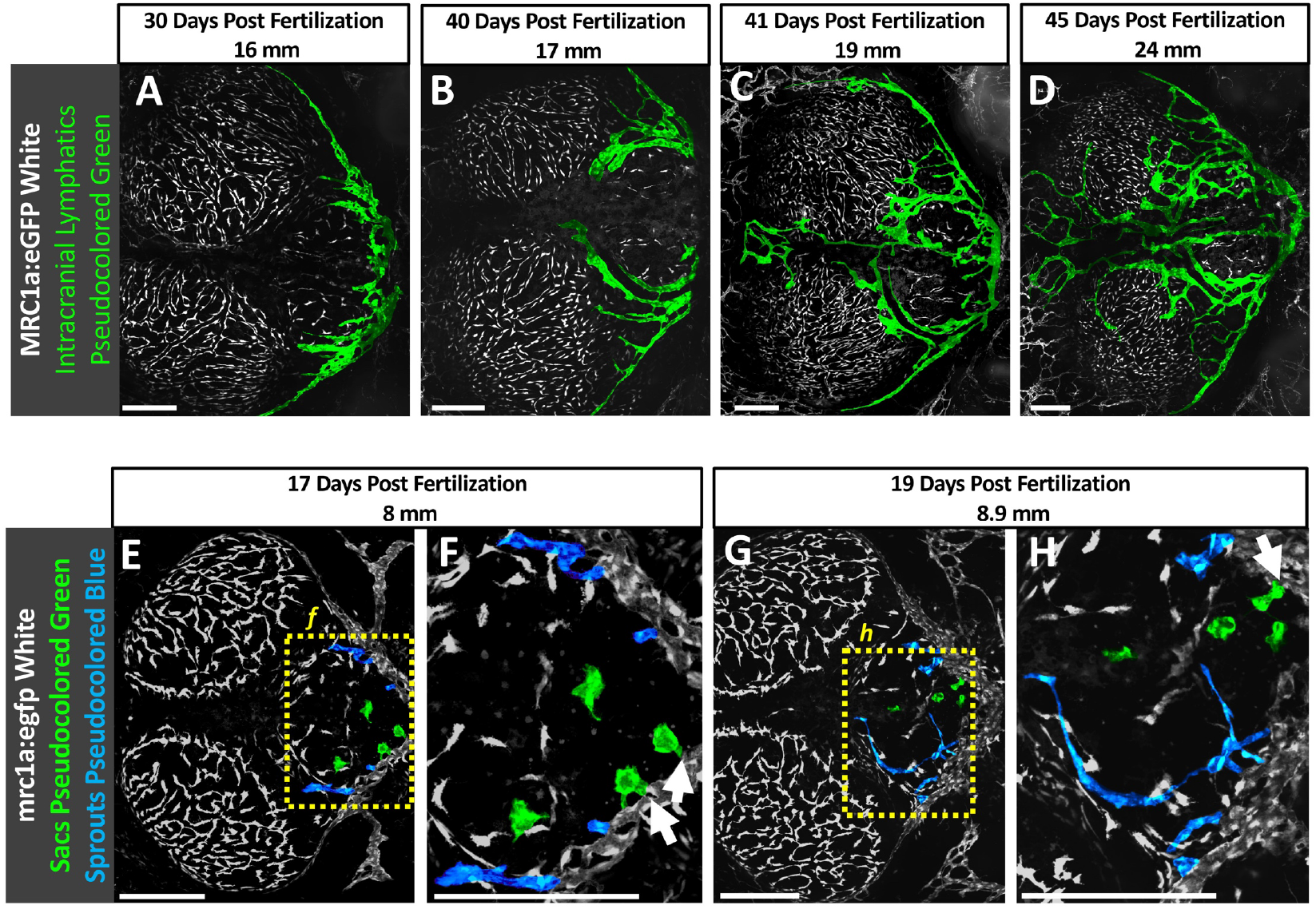
Later development of intracranial lymphatics in juvenile zebrafish. **A-D.** Colored confocal images of mrc1a:egfp+ FGPs (white, uncolored) and intracranial lymphatic vessels (colored green) in the dorsal heads of 30 dpf/16 mm (A), 40 dpf/17 mm (B), 41 dpf/19 mm (C), and 45 dpf/24 mm (D) *Tg(mrc1a:egfp)^y251^* transgenic zebrafish. **E.** Confocal image of mrc1a:egfp+ FGPs and intracranial lymphatic vessels (green) in the dorsal head of a 15 dpf/6.7 mm *Tg(mrc1a:egfp)^y251^* transgenic zebrafish. **F.** Higher magnification view of the boxed region in panel E, showing sprouting intracranial lymphatics (arrows). **G.** Confocal image of mrc1a:egfp+ FGPs and intracranial lymphatic vessels (green) in the dorsal head of a 17 dpf/8 mm *Tg(mrc1a:egfp)^y251^* transgenic zebrafish. **H.** Higher magnification view of the boxed region in panel G, showing presumptive lymphatic sacs fusing with the growing lymphatic vessels (arrows). Scale Bars: 250 um.

### Immune cell trafficking in meningeal lymphatic vessels

In addition to draining interstitial fluid, lymphatic vessels have an important function in trafficking of immune cells. We carried out time-lapse confocal imaging of the intact dorsal heads of living *casper Tg(mrc1a:egfp)^y251^, Tg(lyz:DsRed2)^nz50^* double-transgenic zebrafish to observe whether lyz:dsred+ neurotrophils could be seen transiting through and/or across the walls of mrc1a+ intracranial lymphatics. Live imaging of lymphatic vessels running along the boundary of the cerebellum and optic tectum in an adult *casper* double transgenic animal (**Fig. 7A**) reveals neutrophils moving slowly through the lymphatic vessels (**Fig. 7B-D, Supp. Movie 7**), in contrast to the very rapid movement of neutrophils through nearby blood vessels (**Supp. Movie 7**). In addition to moving through the lymphatics, neutrophils can also occasionally be observed transmigrating through the lymphatic endothelial wall (**Fig. 7E-I, Supp. Movie 8**). Intracranial (**Fig. 7J**) and external superficial (**Fig. 7K**) lymphatic vessels both contain abundant numbers of neutrophils. These data suggest that zebrafish intracranial lymphatics play an important role in immune protection of the brain, as has been suggested for mammalian meningeal lymphatics.

**Fig. 7.**
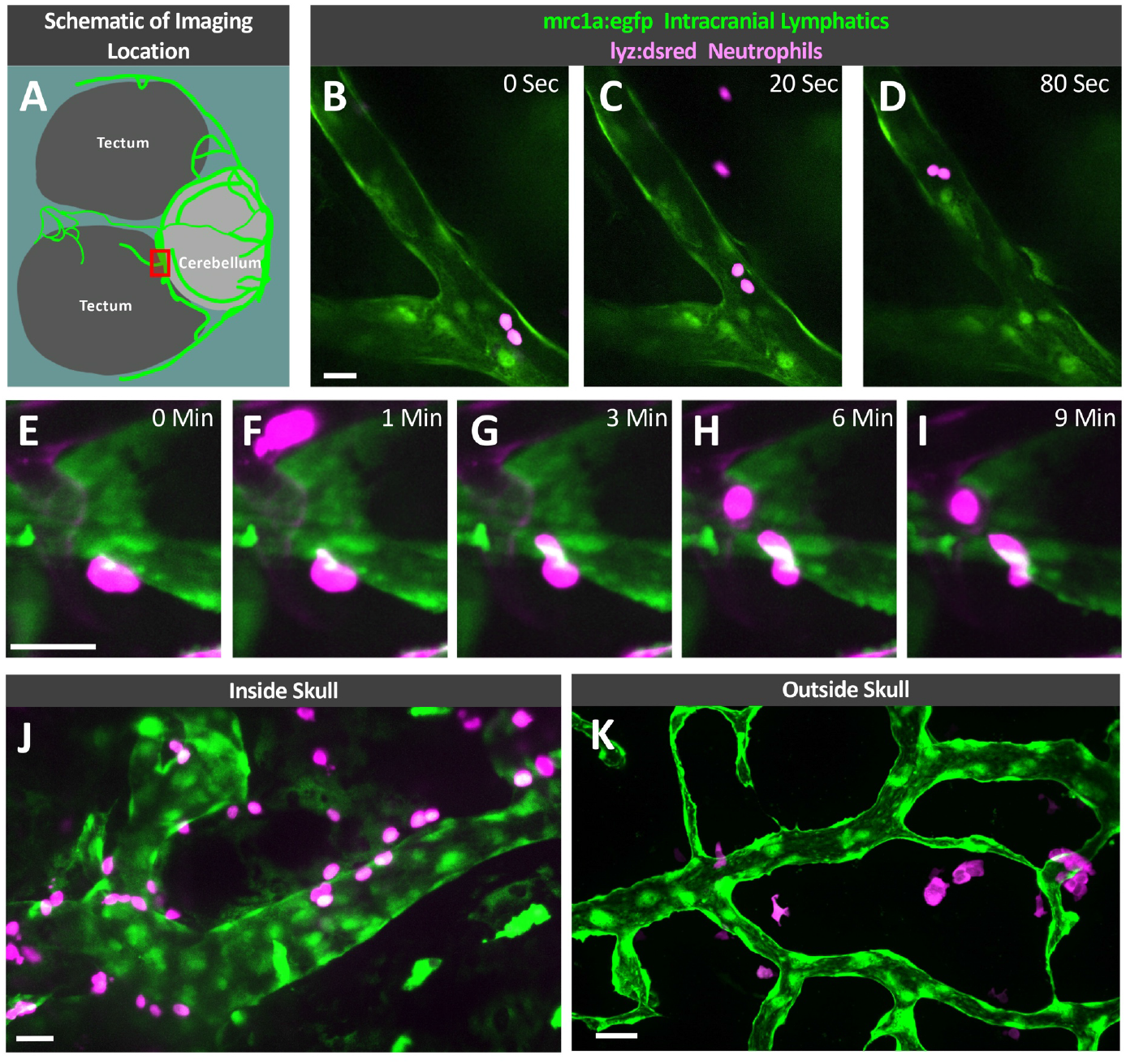
Immune cell trafficking in zebrafish intracranial lymphatics. **A.** Schematic diagram of the optic tecta, cerebellum, and intracranial lymphatics in the dorsal head of an adult *casper, Tg(mrc1a:egfp)^y251^* zebrafish. The red box notes the approximate area shown in the high-magnification images in panels B-D and E-I. **B-D**. Selected frames from a time-lapse confocal image series taken through the skull of a living adult *casper, Tg(mrc1a:egfp)^y251^, Tg(lyz:DsRed2)^nz50^* double transgenic zebrafish showing neutrophils trafficking through an intracranial lymphatic vessel (see **Supp. Movie 7**). **E-I**. Selected frames from a time-lapse confocal image series taken through the skull of a living adult *casper, Tg(mrc1a:egfp)^y251^, Tg(lyz:DsRed2)^nz50^* double transgenic zebrafish showing a neutrophil transmigrating into an intracranial lymphatic vessel (see **Supp. Movie 8**). **J**. Image taken through the skull of a living adult *casper, Tg(mrc1a:egfp)^y251^, Tg(lyz:DsRed2)^nz50^* double transgenic zebrafish showing neutrophils on the inside and outside of an intracranial lymphatic vessel. **K**. Image taken of superficial lymphatics on the dorsal head of a living adult *casper, Tg(mrc1a:egfp)^y251^, Tg(lyz:DsRed2)^nz50^* double transgenic zebrafish showing neutrophils interacting with a lymphatic vessel outside of the skull. Scale bars: 25 um.

## Discussion

In this study we use high-resolution confocal imaging of the meninges in living transgenic zebrafish to show that zebrafish possess a complex and robust meningeal lymphatic network comparable to that found in mammals. We use angiography and intracisternal lymphatic drainage assays to confirm that zebrafish meningeal lymphatics are distinct from the blood vascular network and that they drain interstitial fluid from the brain. We show that intracranial lymphatics initially emerge from the dorsal edge of the previously described facial lymphatic network ^9^, first growing along the caudal edge of the cerebellum and optic tecta before forming a more complex network extending further rostrally. We also show that, like the meningeal lymphatics in mammals, these vessels contain immune cells migrating both through and in and out of them. The discovery of intracranial meningeal lymphatics in the zebrafish provides an important new model for experimental analysis of meningeal lymphatic development and function in health and disease.

The Italian anatomist Paolo Mascagni reported the existence of meningeal lymphatics in humans over 200 years ago ^33^, but his early description was largely ignored and mammalian meningeal lymphatics were only “rediscovered” a few years ago ^1, 2^. It is likely that meningeal lymphatics were overlooked for several centuries at least in part because most researchers were examining isolated brains, and mammalian intracranial lymphatics are found exclusively in the dural layer of the meninges that adheres tightly to the skull when the brain is removed ^1, 2^. Although the dural meninges of the zebrafish have not yet been characterized, we show that, as in mammals, dissected zebrafish brains with the skull removed have no lymphatic vessels associated with them while the inner surface of the zebrafish skull has a complex lymphatic vascular network comparable to that observed in mammals, suggesting that zebrafish intracranial lymphatics may also reside within a dural meningeal layer (**Fig 1**). The elaborate branched network of zebrafish meningeal lymphatics also shares other features with mammalian meningeal lymphatics, including alignment along dural sinuses and drainage of CSF.

We and others recently reported unusual macrophage-like, highly phagocytic cells closely associated with blood vessels in the brain-associated leptomeninges, that despite their macrophage-like morphology display a transcriptional profile highly similar to lymphatic endothelial cells ^18–20^. Although these perivascular cells (called FGPs, Mato cells, muLECs, or BLECs in different reports) have a lymphatic molecular identity, they do not form tubes, at least not under normal physiological conditions, and our findings show that these “lymphatic-like” cells are only present in the brain-associated meninges, unlike the *bona fide* lymphatic vascular network present only in the skull-associated meninges (**Fig 1**). Since FGPs and meningeal lymphatics are actually both external to the brain proper, and since FGPs in particular represent a distinct cell population from vessel-forming lymphatic endothelial cells in meningeal lymphatics, we would suggest that “brain lymphatic endothelial cells” (BLECs) represents a less appropriate designation for Mato cells/FGPs/muLECs. FGPs and meningeal lymphatics also appear to have very distinct functions-FGPs absorb extracellular waste and sequester it in intracellular inclusions, while meningeal lymphatics drain extracellular fluid through an elaborate network of tubes and serve as a route for immune cell trafficking, although future studies may reveal functional interconnections between FGPs and meningeal lymphatics.

By imaging large numbers of animals at different stages of larval and juvenile development we have shown that zebrafish meningeal lymphatics originate from the facial lymphatic plexus prior to skull formation, disconnecting from the facial lymphatics as the skull plates continue to grow and fuse (**Fig. 4,5, Supp. Fig. 4**). In mammals, the first meningeal lymphatic vessels appear at a reasonably comparable anatomical location at the foramen magnum, although in contrast to fish they first begin to appear after the skull has already developed ^34^. Following initial zebrafish intracranial lymphatic network formation at the caudal margins of the cerebellum and optic tecta, further meningeal lymphatic development continues through rostral growth of new lymphatic vessels from this primary plexus. These new lymphatic vessels most often grow alongside major dural blood vessels (**Fig. 6A-D**), with growth apparently taking place both by lymphangiogenic sprouting from the preexisting plexus as well as by formation and incorporation of lymphatic cell clusters or sacs (**Fig. 6E-H**), as has been reported for growing meningeal lymphatics in mice ^34, 35^. In mice, interstitial and cerebrospinal fluid drain through dorsal meningeal lymphatic vessels ^4^ and basal meningeal lymphatic vessels ^3^ emptying into the deep cervical node. Cerebrospinal fluid produced in the choroid plexus flows into the subarachnoid space where it is believed to be drained by arachnoid villi ^36^ and the meningeal lymphatic network ^4, 37^. In addition to draining cerebrospinal fluid and interstitial fluid, the meningeal lymphatic network also drains macromolecules, cellular waste products, and serves as a conduit for trafficking of immune cells ^38^. Our studies suggest that zebrafish meningeal lymphatics also drain interstitial and cerebrospinal fluid and actively traffic immune cells, although due to the challenges of imaging very deep structures in juvenile and adult animals the precise drainage points for this network remain to be determined. At this point we cannot distinguish whether zebrafish intracranial lymphatics drain directly into blood vessels at or near the base of the skull or if they exit the skull and connect to other lymphatic vessels. The possibility that the dorsal meningeal lymphatic network of the zebrafish drains into an as-yet-undiscovered structure analogous to the deep cervical lymph node also cannot be ruled out.

The existence of a meningeal lymphatic network in the zebrafish provides a unique and useful new model for visualization and for experimental and genetic manipulation of this critical vascular system. The thin skull of the zebrafish permits high resolution optical imaging of meningeal lymphatics in living adult animals, while the vast array of transgenic tools, mutants, and disease models available in the fish facilitates investigation of the development, anatomy, and function of this system. As noted in the introduction, exciting new research has suggested that intracranial lymphatics play important roles in and/or may provide an important gateway for treatment of a variety of brain pathologies including Alzheimer’s disease ^5, 37^, brain cancer ^6, 39, 40^, multiple sclerosis ^4^, and meningitis. Zebrafish models of meningitis have in fact recently been developed ^41, 42^ and examining the role of meningeal lymphatic vessels in these models should prove to be a fruitful avenue of study. Overall, the ability to carry out high resolution optical imaging and experimental and genetic manipulation in the fish, and to maintain large number of fish and to obtain enormous numbers of their embryonic and larval progeny for analysis, make this a superb model to pursue large scale genetic and chemical screens, as well as other future studies aimed at understanding more about this important but heretofore elusive vascular system and how it functions in health and disease.

## Acknowledgments

The authors would like to thank members of the Weinstein laboratory for their critical comments on this manuscript. The Authors would also like to thank the Research Animal Branch of the *Eunice Kennedy Shriver* National Institute of Child Health and Human Development as well as the Charles River Staff for excellent animal care and husbandry.

## Supplemental Materials

Online Figures 1-4

Online Videos 1-8

## Non-standard Abbreviations and Acronyms

VEGF-C: Vascular Endothelial Growth Factor C
FGPs: Fluorescent Granular Perithelial cells
muLECs: meningeal mural Lymphatic Endothelial Cells
BLECs: Brain Lymphatic Endothelial cells
CSF: cerebrospinal fluid
mrc1a: mannose receptor, C type 1a
lyve1b: lymphatic vessel endothelial hyaluronic receptor 1b
dpf: Days Post Fertilization

