## Supplemental Figs. 1-4 for "Live Imaging of Intracranial Lymphatics in the Zebrafish"

### SUPPLEMENTAL MATERIALS

### SUPPLEMENTAL FIGURES

**Supplemental Figure 1, Castranova et al.**

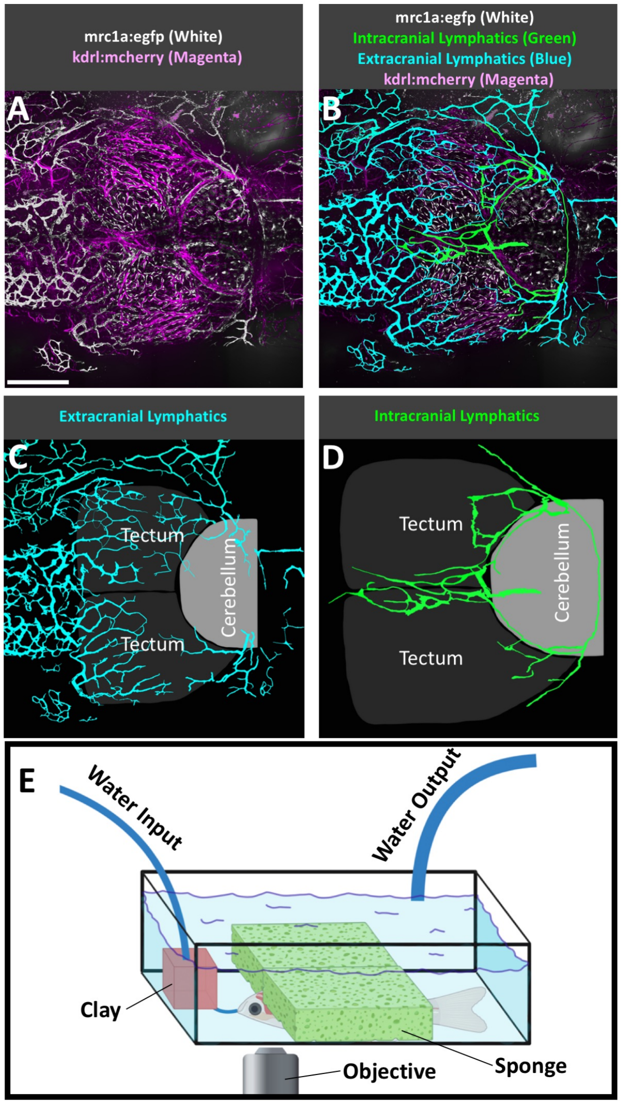


**Supp. Fig. 1 Superficial lymphatic networks on the adult zebrafish head**

**A**. Confocal dorsal view image of an adult *casper*, *Tg(mrc1a:egfp)*$^{y251}$, *Tg(kdrl:mcherry)*$^{y206}$ double transgenic zebrafish head, showing kdrl:mcherry+ blood vessels in magenta and mrc1a:egfp+ superficial lymphatics, intracranial lymphatics, and FGPs in white. Rostral is to the left. **B**. Colored version of panel A highlighting the superficial lymphatic network in teal and the intracranial lymphatic network in green. Blood vessels and FGPs are shown in their original colors of magenta and white respectively. **C**. Colored version of the mrc1a:egfp+ vessels from panel A, showing only the superficial (outside the skull) lymphatic network (in teal) in relation to optic tecta and cerebellum. **D**. Colored version of the mrc1a:egfp+ vessels from panel A, showing only the intracranial (inside the skull) lymphatic network (in green) in relation to the optic tecta and cerebellum. **E**. Schematic of zebrafish intubation method we designed for longer term imaging of juvenile or adult zebrafish on an inverted microscope, adapted from Xu et al.[25] Scale bar: 500 um.

**Supp. Figure 2, Castranova et al.**

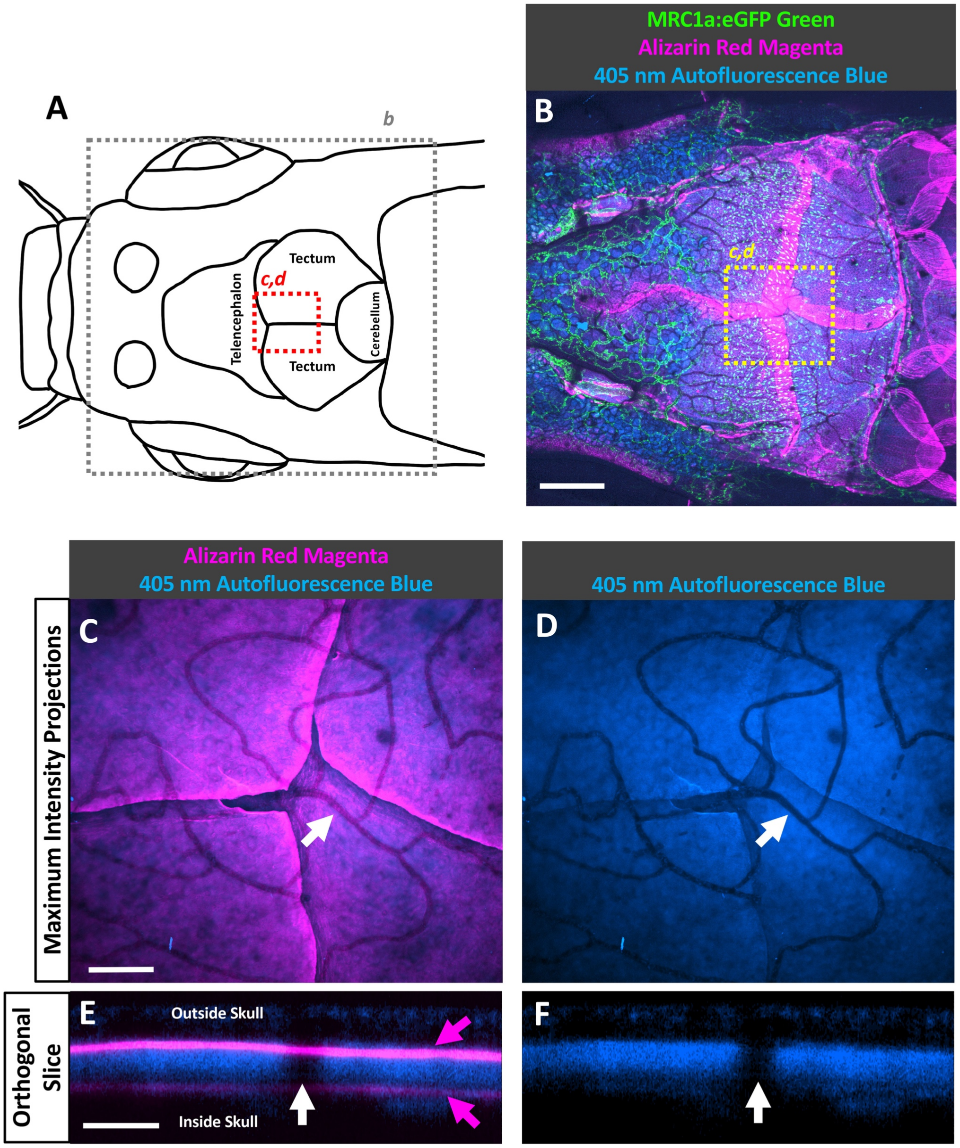


**Supp. Fig. 2 The zebrafish skull autofluoresces under violet light (405 nm)**

**A.** Schematic dorsal view diagram of an adult zebrafish head showing the telencephalon, optic tecta, and cerebellum, with the approximate regions imaged in panel B shown with a grey box and in panels C and D shown with a red box. **B.** Confocal dorsal view image of a 25 mm adult *casper*, *Tg(mrc1a:egfp)*$^{y251}$ transgenic zebrafish head stained with Alizarin red to show calcified bone, and also showing autofluorescence produced by 405 nm laser excitation. The yellow box notes the region shown in the higher-magnification images in panels C and D. **C,D.** Higher magnification alizarin+ skull autofluorescence (C) or skull autofluorescence alone (D) images of the yellow boxed region in panel B, showing overlapping skull plates. The white arrows indicate areas of strongly reduced 405 nm autofluorescence where blood vessels are present inside the skull. **E,F.** Orthogonal slices from small areas of panels C and D, respectively, showing the same area (white arrows) where blood vessels are present inside the skull creating a gap in autofluorescence. Magenta arrows show Alizarin red fluorescence is limited to a thin layer inside and outside the skull, with the outside signal being significantly stronger, while autofluorescence, though weaker, is detected throughout the entire thickness of the skull. Scale bars: 500 um (B), 100 um (C), 20 um (D).

### Supp Fig Figure 3, Castranova et al.

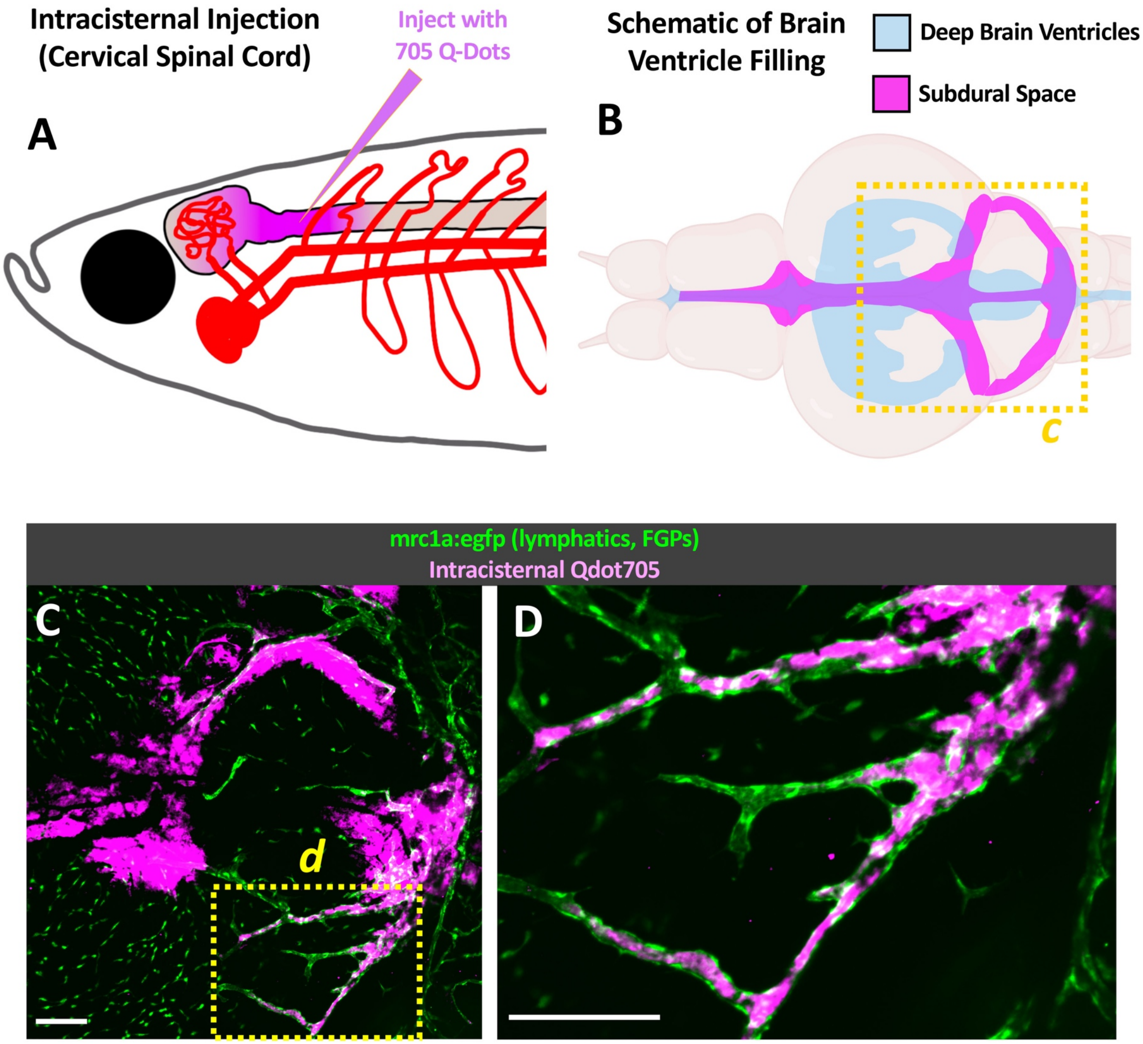


**Supp. Fig. 3 Intracisternal spinal injections drain into intracranial lymphatics**

**A.** Lateral view schematic diagram showing the injection site of Q-Dot705 into the cervical spinal cord of a one month old *casper*, *Tg(mrc1a:egfp)*$^{y251}$ transgenic zebrafish. **B.** Dorsal view schematic diagram showing areas within and surrounding the zebrafish brain filled by spinally injected quantum dots, including the deep brain ventricles (light blue) and the subdural space (magenta) with the approximate area imaged in C shown by the dashed box. **C**. Confocal imaging of the dorsal head of a one month old *casper*, *Tg(mrc1a:egfp)*$^{y251}$ transgenic zebrafish, showing Q-dot705 injected into the cervical spinal cord draining into the subdural space and being collected by lymphatic vessels. The yellow dashed box notes the region shown in the higher magnification image in panel D. **D**. Higher magnification image of the boxed region from panel C, showing Q-dot705 filled lymphatics. Scale bars: 150 um.

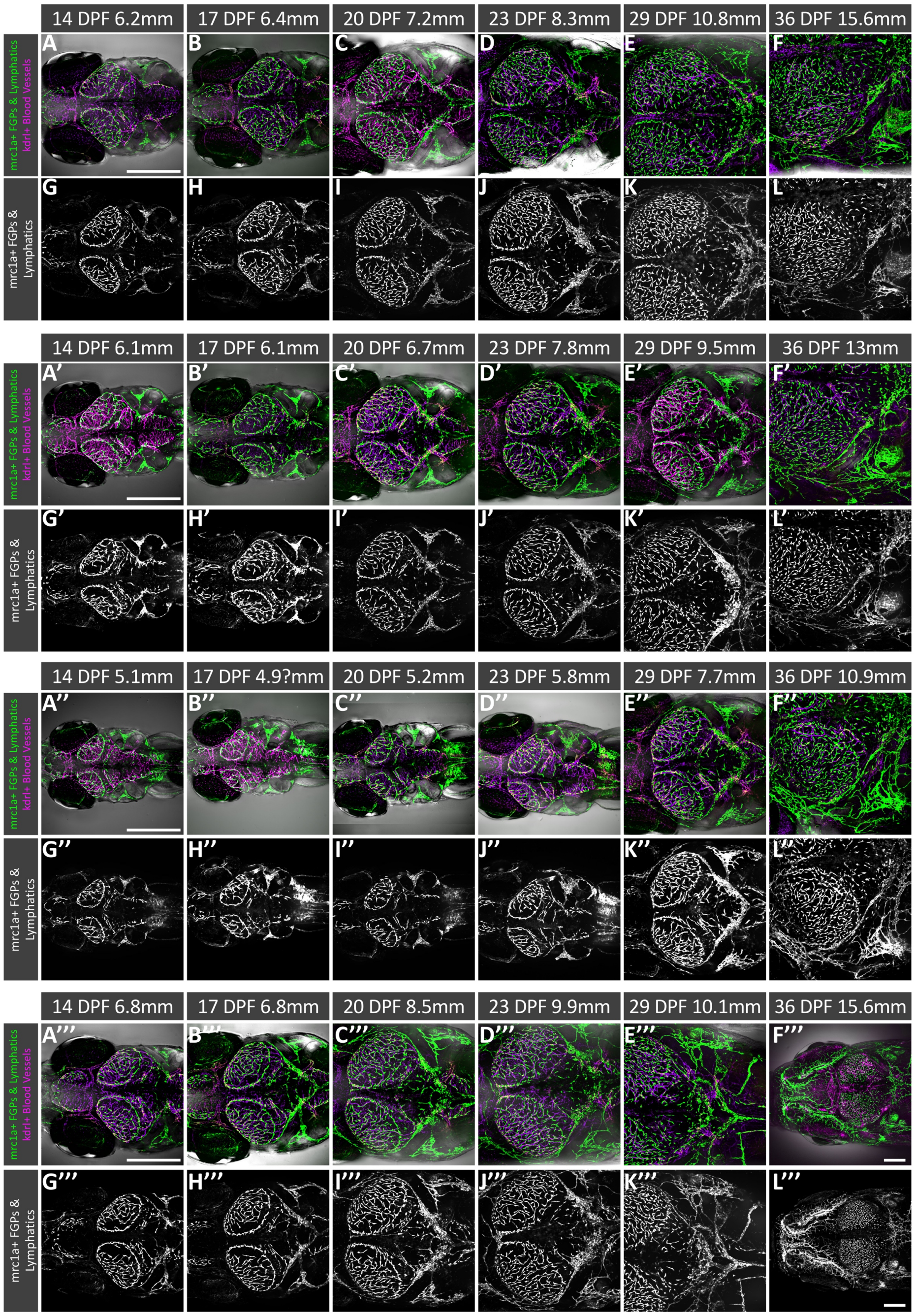


**Supp Fig. 4** **Time series of initial intracranial lymphatic network development**

**A-L.** Consecutive dorsal view confocal images of the head of the same adult *casper*, *Tg(mrc1a:egfp)*$^{y251}$, *Tg(kdrl:mcherry)*$^{y206}$ double transgenic zebrafish ("Fish #1") collected at 14 dpf (A,G), 17 dpf (B,H), 20 dpf (C,I), 23 dpf (D,J), 29 dpf (E,K), and 36 dpf (F,L). **A-F.** Composite images showing mrc1a:egfp+ lymphatic vessels (green), kdrl:mcherry+ blood vessels (magenta) and DIC transmitted light images (greyscale, displayed as focus stacked projections). **G-L.** Maximum intensity greyscale image of mrc1a:egfp+ lymphatics and FGPs. **A'-L'.** As for A-L, but images are of Fish #2. **A"-L".** As for A-L, but images are of Fish #3. **A'"-L'".** As for A-L, but images are of Fish #4. The lengths of each fish measured at the time that each image stack was collected are noted above each image. Scale bars: 500 um.

### SUPPLEMENTAL MOVIE LEGENDS

#### Supp. Movie 1 3D rotation of adult intracranial lymphatics

3D visualization of a confocal Z-stack of the dorsal head of an adult *casper*, *Tg(mrc1a:egfp)*$^{y251}$, *Tg(kdrl:mcherry)*$^{y206}$ double transgenic zebrafish, showing mrc1a:egfp+ intracranial lymphatics and FGPs (green), kdrl:mcherry+ blood vessels (magenta), and 405 nm skull autofluorescence (blue).

#### Supp. Movie 2 Adult intracranial lymphatics Z-stack scroll-through

Scroll through a confocal Z-stack of the dorsal head of an adult *casper*, *Tg(mrc1a:egfp)*$^{y251}$, *Tg(kdrl:mcherry)*$^{y206}$ double transgenic zebrafish, showing mrc1a:egfp+ intracranial lymphatics and FGPs (green), kdrl:mcherry+ blood vessels (magenta), and 405 nm skull autofluorescence (blue).

#### Supp. Movie 3 Intracranial lymphatic drainage

3D visualization and scroll-through a confocal Z-stack of he dorsal head of an adult *casper*, *Tg(mrc1a:egfp)*$^{y251}$, *Tg(kdrl:mcherry)*$^{y206}$ double transgenic zebrafish injected intracranially with Qdot705 and blue dextran, showing mrc1a:egfp+ lymphatics (white), kdrl:mcherry+ blood vessels (yellow), 405 nm skull autofluorescence (blue), and intralymphatic 705 nm Qdot705 fluorescence (magenta) and blue dextran fluorescence (blue). The 3D rotation and scroll through are followed by a 25x sped up real-time movie sequence of a single plane containing an intracranial blood vessel (yellow) next to an intracranial lymphatic vessel (white) filled with Qdot705 (magenta) and blue dextran (blue). The movie shows the same vessels as in **Figure 3F-K**.

#### Supp. Movie 4 Primary intracranial lymphatic network formation

3D visualization of confocal Z-stacks of the dorsal head of adult *casper*, *Tg(mrc1a:egfp)*$^{y251}$, *Tg(kdrl:mcherry)*$^{y206}$ double transgenic zebrafish at 9 dpf (4mm), 17 dpf (6mm), 23 dpf (7.8 mm), and 27 dpf (10.3 mm), showing mrc1a:egfp+ lymphatics and FGPs (green) and kdrl:mcherry+ blood vessels (magenta). Orange and blue coloring at various points in the movie sequence highlights the developing superficial and intracranial lymphatics, respectively. The movie is showing the same image stacks depicted in **Figure 4**.

#### Supp. Movie 5 Intracranial lymphatics and the skull at 17 dpf

3D visualization of a confocal Z-stack of the dorsal head of a 17 dpf (8.2 mm) *casper*, *Tg(mrc1a:egfp)*$^{y251}$, *Tg(Ola.Sp7:mCherry-Eco.NfsB)*$^{pd46}$ double-transgenic zebrafish, showing mrc1a:egfp+ lymphatic vessels and FGPs (green) and sp7:mcherry+ developing skull plates (magenta). Orange and blue coloring at a later point in the movie sequence highlights developing superficial facial and intracranial lymphatics, respectively.

#### Supp. Movie 6 Intracranial lymphatics and the skull 19 dpf

3D visualization of a confocal Z-stack of the dorsal head of a 19 dpf (8.9 mm) *casper*, *Tg(mrc1a:egfp)*$^{y251}$, *Tg(Ola.Sp7:mCherry-Eco.NfsB)*$^{pd46}$ double-transgenic zebrafish, showing mrc1a:egfp+ lymphatic vessels and FGPs (green) and sp7:mcherry+ developing skull plates (magenta). Orange and blue coloring at a later point in the movie sequence highlights developing superficial facial and intracranial lymphatics, respectively. The movie is showing the same image stack depicted in **Figure 5F-H**.

#### Supp. Movie 7 Neutrophil trafficking

Real-time confocal image series of a single Z plane taken through the dorsal head of a living adult *casper*, *Tg(mrc1a:egfp)$^{y251}$, Tg(lyz:DsRed2)$^{nz50}$* double transgenic zebrafish showing neutrophils (magenta; lyz:dsred+) trafficking through an intracranial lymphatic vessel. The movie is showing the same image stack depicted in **Figure 7B-D**.

#### Supp. Movie 8 Neutrophil transmigration

3D visualization of confocal Z-stacks of the dorsal head of a living adult *casper*, *Tg(mrc1a:egfp)$^{y251}$* (green)*, Tg(lyz:DsRed2)$^{nz50}$* (magenta) *Tg(kdrl:mcherry)$^{y206}$* (magenta) triple transgenic zebrafish, followed by a time-lapse confocal image series of a single Z plane showing a neutrophil transmigrating into an intracranial lymphatic vessel. The movie is showing the same image stack depicted in **Figure 7E-I**.
